# Species limits and hybridization in Andean leaf-eared mice (*Phyllotis*)

**DOI:** 10.1101/2024.08.31.610610

**Authors:** Marcial Quiroga-Carmona, Schuyler Liphardt, Naim M. Bautista, Pablo Jayat, Pablo Teta, Jason L. Malaney, Tabitha McFarland, Joseph A. Cook, Moritz L. Blumer, Nathanael D. Herrera, Zachary A. Cheviron, Jeffrey M. Good, Guillermo D’Elía, Jay F. Storz

**Affiliations:** School of Biological Sciences, University of Nebraska, Lincoln, NE, United States; Instituto de Ciencias Ambientales y Evolutivas, Facultad de Ciencias, Universidad Austral de Chile, Valdivia, Chile; Colección de Mamíferos, Facultad de Ciencias, Universidad Austral de Chile, Campus Isla Teja, Valdivia, Chile; Division of Biological Sciences, University of Montana, Missoula, MT, United States; Unidad Ejecutora Lillo (CONICET-Fundación Miguel Lillo), San Miguel de Tucumán, Argentina; Departamento de Ciencias Básicas y Tecnológicas, Universidad Nacional de Chilecito (UNdeC), Argentina; División Mastozoología, Museo Argentino de Ciencias Naturales “Bernardino Rivadavia”, Ciudad Autónoma de Buenos Aires, Argentina; New Mexico Museum of Natural History and Science, Albuquerque, NM, United States; Museum of Southwestern Biology, University of New Mexico, Albuquerque, NM, United States; Department of Biology, University of New Mexico, Albuquerque, NM, United States; Department of Genetics, University of Cambridge, Cambridge, United Kingdom

**Keywords:** Altiplano, Andes, geographic range limits, introgression, Phyllotini, Sigmodontinae, species delimitation

## Abstract

Leaf-eared mice (genus *Phyllotis*) are among the most widespread and abundant small mammals in the Andean Altiplano, but species boundaries and distributional limits are often poorly delineated due to sparse survey data from remote mountains and high-elevation deserts. Here we report a combined analysis of mitochondrial DNA variation and whole-genome sequence (WGS) variation in *Phyllotis* mice to delimit species boundaries, to assess the timescale of diversification of the group, and to examine evidence for interspecific hybridization. Estimates of divergence dates suggest that most diversification of *Phyllotis* occurred during the past 3 million years. Consistent with the Pleistocene Aridification hypothesis, our results suggest that diversification of *Phyllotis* largely coincided with climatically induced environmental changes in the mid- to late Pleistocene. Contrary to the Montane Uplift hypothesis, most diversification in the group occurred well after the major phase of uplift of the Central Andean Plateau. Species delimitation analyses revealed surprising patterns of cryptic diversity within several nominal forms, suggesting the presence of much undescribed alpha diversity in the genus. Results of genomic analyses revealed evidence of ongoing hybridization between the sister species *Phyllotis limatus* and *P. vaccarum* and suggest that the contemporary zone of range overlap between the two species represents an active hybrid zone.

## 1. INTRODUCTION

Leaf-eared mice in the genus *Phyllotis*, Waterhouse 1873, are emblematic mammals of the Andean Altiplano and have an exceptionally broad latitudinal distribution in South America, from Ecuador to the northern coast of the Strait of Magellan (Steppan and Ramírez, 2015). The genus has an even more impressive elevational distribution: Whereas *P. darwini* is found at sea level along the desert coastline of northern Chile, and species like *P*. *anitae*, *P*. *nogalaris*, and *P*. *osilae* are found in humid, lowland Yungas forests on the eastern sub-Andean slopes (Jayat et al., 2016), other taxa such as *P. vaccarum* have been documented at extreme elevations (>6000 m above sea level) on the upper reaches and summits of some of the highest peaks in the Andean Cordillera (Storz et al., 2020, 2023, 2024; Steppan et al., 2022). Although *Phyllotis* mice are among the most widespread and abundant small mammals in the Andean Altiplano and adjacent lowlands, the taxonomic status and range limits of many species are not well-resolved due to sparse survey data from remote mountains and high-elevation deserts (puna). The resultant gaps in sampling coverage have hindered a complete assessment of species richness and geographic distributions of *Phyllotis* mice.

Over the last two decades, *Phyllotis* has been subject to several taxonomic assessments that have helped resolve species limits and phylogenetic relationships (Jayat et al., 2007, 2016, 2021; Ojeda et al., 2021; Steppan et al., 2007; Rengifo and Pacheco, 2015, 2017; Teta et al., 2018, 2022). There are currently 26 recognized species of *Phyllotis*, and the genus comprises three main clades, commonly referred to as the *andium-amicus*, *osilae*, and *darwini* species groups (Rengifo and Pacheco, 2017; Steppan, 1993, 1995; Steppan et al., 2007; Steppan and Ramírez, 2015; Teta et al., 2022). The *darwini* group is the most speciose and includes several species that are distributed in the Atacama Desert and the Andean dry puna: *P. caprinus*, ‘*P. chilensis-posticalis*’ (*sensu* Pearson, 1958; referred to as ‘*P*. *posticalis*-*rupestris*’ by Ojeda et al., 2021), *P. darwini*, *P*. *limatus*, *P*. *magister*, *P*. *osgoodi*, and *P. vaccarum* (Jayat et al., 2021; Ojeda et al., 2021; Steppan and Ramírez, 2015; Teta et al., 2022; Storz et al., 2024). In northeastern Chile and bordering regions of Argentina, Bolivia, and Peru, the ranges of several of these species potentially overlap (Fig. 1A), but in most cases the distribution limits are not clearly defined. We often do not know the extent to which species ranges overlap across Andean elevational gradients, which is important for understanding the relative roles of competitive exclusion and physiological tolerances in shaping elevational patterns of species turnover and for detecting distributional shifts in response to climate change.

**Figure 1.**
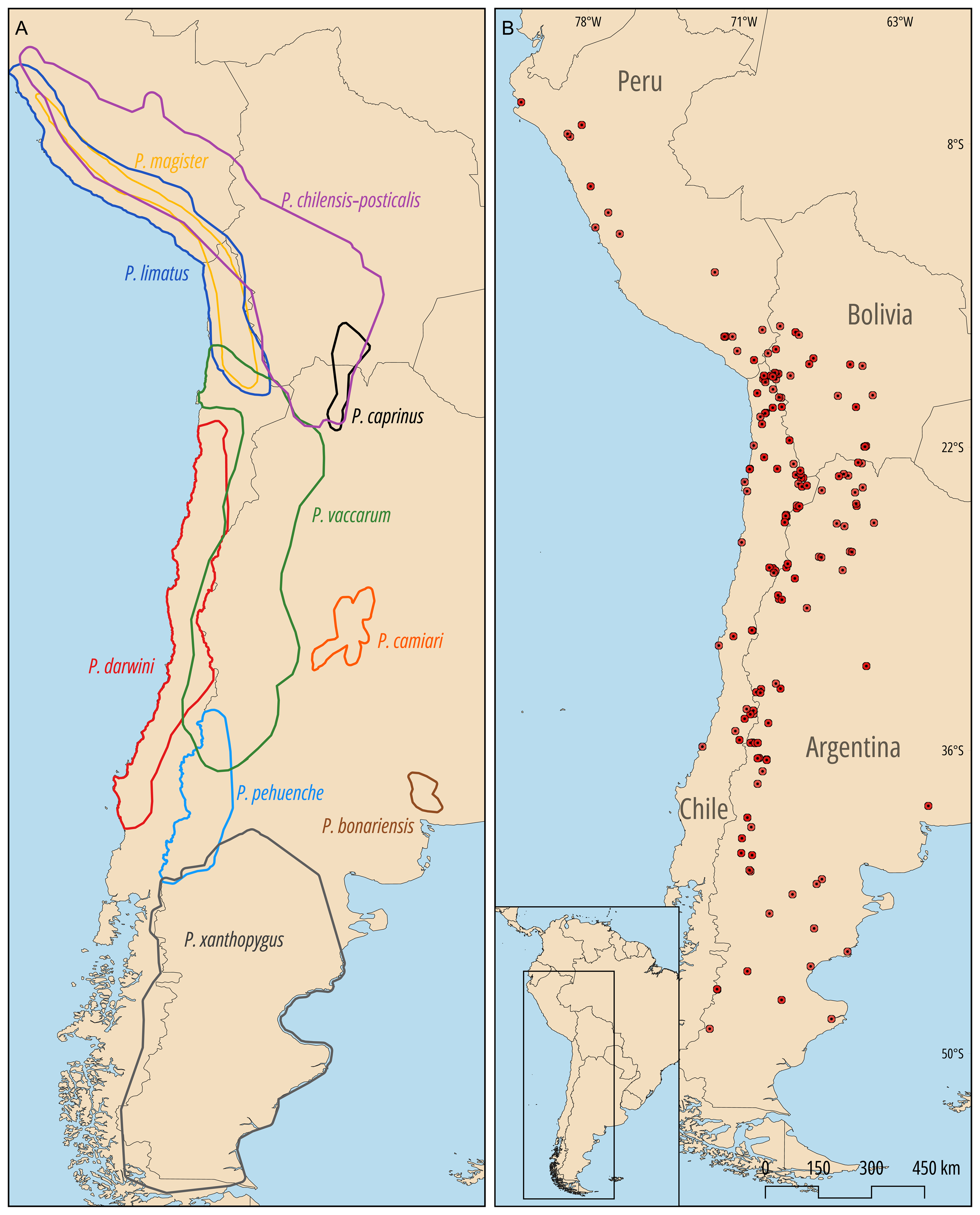
Distribution limits of *Phyllotis* species and geographic sampling coverage in the Central Andes and adjoining lowlands. A) Ranges of *Phyllotis* mice in the *P*. *darwini* species group, based on patterns of morphological and DNA marker variation (Jayat et al., 2021; Ojeda et al., 2021; Steppan and Ramírez, 2015; Storz et al., 2024). B) Distribution of 169 sampling localities, representing sites of origin for 448 *Phyllotis* specimens used in the survey of *cytb* and WGS variation.

In this high-Andean region, genomic delimitation of species boundaries between *P. limatus* and *P. vaccarum* in northern Chile led to a dramatically revised understanding of the latitudinal and elevational range limits of the former species (Storz et al., 2024). Previously inferred range limits of *P. limatus* were found to be in error because specimens from the highest elevations and most southern latitudes had been mis-identified as *P. limatus* on the basis of mitochondrial (mt) DNA and were later that some *P. vaccarum* carry mtDNA haplotypes more closely related to those of *P. limatus* suggests a history of introgressive hybridization and/or incomplete lineage sorting. In addition to highlighting the importance of using multilocus data to delimit species boundaries, the observed mitonuclear discordance between *P. limatus* and *P. vaccarum* suggests the possibility of hybridization between other pairs of species of *Phyllotis* in regions of historical or contemporary range overlap.

Here we report a combined analysis of mtDNA variation and whole-genome sequence (WGS) variation in mice of the genus *Phyllotis* aimed to delimit species boundaries, to assess the timescale of diversification of the group, and to examine evidence for interspecific hybridization. The analysis is principally focused on a large set of vouchered specimens that we collected over the course of five high-elevation survey expeditions in the Puna de Atacama, Central Andes (2020-2023), in conjunction with additional collecting trips in the surrounding Altiplano and adjoining lowlands in Argentina, Bolivia, and Chile. The genomic analysis is primarily focused on members of the *P*. *darwini* species group that have overlapping or potentially overlapping ranges.

## 2. MATERIAL AND METHODS

### 2.1 Specimen collection

We collected representatives of multiple species of *Phyllotis* during the course of small-mammal surveys in the Altiplano and adjoining lowlands on both sides of the Andean Cordillera in Chile, Bolivia, and Argentina. We captured all mice using Sherman live traps, in combination with Museum Special snap traps in some localities. We sacrificed animals in the field, prepared them as museum specimens, and preserved liver tissue in ethanol as a source of genomic DNA. All specimens are housed at Colección de Mamíferos de la Universidad Austral de Chile, Valdivia, Chile (UACH), Colección Boliviana de Fauna, La Paz, Bolivia (CBF), Centro Regional de Investigaciones Científicas y Transferencia Tecnológica de La Rioja, La Rioja, Argentina (CRILAR), Centro Nacional Patagónico, Chubut, Argentina (CNP), Fundación-Instituto Miguel Lillo, Tucumán, Argentina (CML), Museo Argentino de Ciencias Naturales “Bernardino Rivadavia”, Ciudad Autónoma de Buenos Aires, Argentina (MACN-Ma), and Museum of Southwestern Biology, New Mexico, USA (MSB). We identified all specimens to the species level based on external characters (Jayat et al., 2021; Steppan and Ramírez, 2015; Teta et al., 2022) and, as described below, we later confirmed field-identifications with DNA sequence data.

In Chile, all animals were collected in accordance with permissions to JFS, MQC, and GD from the following Chilean government agencies: Servicio Agrícola y Ganadero (6633/2020, 2373/2021, 5799/2021, 3204/2022, 3565/2022, 911/2023 and 7736/2023), Corporación Nacional Forestal (171219, 1501221, and 31362839), and Dirección Nacional de Fronteras y Límites del Estado in accordance with permissions to JFS (Resolución Administrativa 026/09) and JAC (DVS-CRT-02/91) from the Ministerio de Medio Ambiente y Agua, Estado Plurinacional de Bolivia. In Argentina, all animals were collected in accordance with the following permissions to JPJ from the Secretaria de Ambiente, Ministerio de Produccion y Ambiente de La Rioja (Expte. N° P4-00402-21 Disp. S.A. N° 001/22 and Expte. N° P4 -00158 -22 Disp. S.A. N° 007/22), the Ministerio de Ambiente, Secretaria de Biodiversidad y Desarrollo Sustentable de Jujuy (Expte. N° 1102-122-2020/SByDS), and the Ministerio de Desarrollo Productivo, Direccion de Flora, Fauna Silvestre y Suelos de Tucumán (Expte. N° 677- 330-2021). All live-trapped animals were handled in accordance with protocols approved by the Institutional Animal Care and Use Committee (IACUC) of the University of Nebraska (project ID’s: 1919, 2100), IACUC of the University of New Mexico (project ID’s: 16787 and 20405), and the bioethics committee of the Universidad Austral de Chile (certificate 456/2022).

### 2.2 sequence data

To maximize geographic coverage in our survey of mtDNA variation, we generated sequence data for a subset of our own voucher specimens (*n*=269) and supplemented this dataset with publicly available *Phyllotis* sequences from GenBank (*n*=179). This sequence dataset (Supplementary Table 1), based on a total of 448 specimens, includes 20 of the 26 nominal species that are currently recognized within the genus *Phyllotis*. We used a subset of our newly collected voucher specimens (*n*=137) for the analysis of WGS variation.

### 2.3 Mitochondrial DNA variation

For the analysis of mtDNA variation, we extracted DNA from liver samples and PCR-amplified the first 801 base pairs of the *cytochrome b* (*cytb*) gene using the primers MVZ 05 and MVZ 16 (Smith and Patton 1993), following protocols of Cadenillas and D’Elía (2021). Of the 269 *cytb* sequences that we generated from our own set of voucher specimens, 89 were published previously (Storz et al., 2020, 2024; GenBank accession numbers: OR784643-OR784661, OR799565-OR799614, and OR810731- OR810743). We deposited all newly generated sequences in GenBank (accession numbers: PQ295377-PQ295555). The newly generated sequences derive from voucher specimens housed in the Argentine, Bolivian, Chilean, and US collections mentioned above (section 2.1).

### 2.4 phylogeny estimation

As outgroups for the phylogenetic analysis, we used *cytb* sequences from five other phyllotine rodents (*Auliscomys boliviensis*, JQ434420; *A*. *pictus*, U03545; *A*. *sublimis*, U03545; *Calomys musculinus*, HM167822; and *Loxondontomys micropus*, GU553838). The final set of 453 sequences was aligned with MAFFT v7 (Katoh et al., 2017) using the E-INS-i strategy to establish character primary the presence of internal stop codons and shifts in the reading frame. Pairwise genetic distances and their standard errors (p-dist./SE) were calculated using MEGA X 10.1.8 (Kumar et al., 2018).

Redundant *cytb* sequences were identified and discarded using the functions *FindHaplo* and *haplotype* in the *sidier* (Pajares, 2013) and *haplotypes* (Aktas, 2023) R packages, respectively. The final matrix of nonredundant sequences included a total of 287 haplotypes.

The nucleotide substitution model (HKY + I + G) that provided the best fit to the nonredundant *cytb* data matrix was selected based on the Bayesian Information Criterion (BIC) using ModelFinder (Kalyaanamoorthy et al., 2017). Genealogical relationships among haplotypes of *Phyllotis* species were estimated via Maximum Likelihood (ML) and Bayesian Inference (BI). The ML analysis was performed using IQ-TREE (Trifinopoulos et al., 2016), with perturbation strength set to 0.5 and the number of unsuccessful iterations set to 100. Nodal support was assessed through 1000 ultrafast bootstrap replicates (UF; Minh et al., 2013). BI was implemented with BEAST 2 v2.6.7 (Bouckaert et al., 2014), which was also used to estimate divergence dates among *Phyllotis* species. A gamma site model was selected with the substitution model set to HKY. The gamma shape parameter (exponential prior, mean 1.0) and proportion of invariant sites (uniform distribution, 0.001–0.999, lower and upper bounds) were estimated. To prevent the sampling of excessively small values for the HKY exchangeability rates, the prior sampling distribution was set to gamma with a shape parameter (alpha) of 2.0 and a scale parameter (beta) of 0.5. The clock model was set to Relaxed Log Normal with an estimated clock rate. The calibrated Yule model was selected to parameterize fossil calibrations. For the mean branch rate (ucldMean), an exponential sampling distribution was applied with a mean of 10.0 and no offset. Given that variation in substitution rates among branches is low and evidence suggests that molecular evolution is largely clock-like across Phyllotini (Parada et al., 2013), standard deviation in rates across branches (ucldStdev) was converted to an exponential prior distribution with a mean of 0.3337 and no offset. Since the fossil record for *Phyllotis* is not sufficient to establish primary calibration points (Pardiñas et al., 2002), we used secondary calibration points from a phylogenetic analysis of the subfamily Sigmodontinae (Parada et al., 2015). We used normal distributions and 95% credibility intervals for estimated crown ages of the genus *Phyllotis* (3.35-6.66 Mya), and the *darwini* species group (4.51-1.77 Mya). We performed two runs of 600 x 10^6^ MCMC generations with trees sampled every 4 x 10^3^ steps, yielding 15,001 samples for parameter estimates. Effective sample sizes greater than 200 for all parameters (i.e., stable values of convergence) were verified using Tracer v1.7.1 (Rambaut et al., 2018). Runs were combined with LogCombiner v2.6.7 (Bouckaert et al., 2014), using a 10% burn-in that was determined by examining individual traces. The first 10% of estimated trees were discarded and the remainder were used to construct a maximum clade credibility tree with posteriori probability values (PP) and age estimates employing TreeAnnotator v2.6.2 (Rambaut and Drummond, 2019).

### 2.5 Assessment of species limits within the *P*. *xanthopygus* species complex

To delimit species within the *P*. *darwini* group, we employed the Bayesian time calibrated-ultrametric tree estimated with BEAST 2 and two single-locus coalescent methods: The General Mixed Yule Coalescent model (GMYC; Pons et al., 2006; Fujisawa and Barraclough, 2013) and the Poisson Tree Processes (PTP; Zhang et al., 2013). Both methods are based on the fit of different mixed models (the General Mixed Yule Coalescent model in the case of the GMYC, and the Poisson Tree Processes in the case of the PTP) to processes of interspecific diversification and/or genealogical branching within species (Fujisawa and Barraclough, 2013; Zhang et al., 2013). These methods were implemented via their online web servers: https://species.h-its.org/gmyc/ and http://species.h-its.org/ptp/, respectively. The Bayesian implementations of these methods (b-GMYC: Reid and Carstens, 2012; b-PTP: Zhang et al., 2013) were also employed to account for uncertainty in gene tree estimation. The b-GMCY analysis was implemented in R via the *b-GMCY* R package (Reid and Carstens 2012), which offers estimates of the posterior marginal probabilities for candidate species, setting a post-burn-in sample of 1000 trees sampled from the posterior distribution of trees. For all parameters, priors were set as default (i.e., t1 and t2 were set at 2 and 100, respectively), and the analysis was completed with 50 x 10^3^ generations, burning 10% of these and with a thinning interval of 1000 samples. The b-PTP analysis was implemented in the associated online web server (http://species.h-its.org/b-ptp/) with default values (i.e., 100 x 10^3^ MCMC, thinning of 100 and burning of 0.1). Branch lengths are proportional to coalescence times in the GMYC model, whereas they are proportional to the number of nucleotide substitutions in the PTP model (Dellicour and Flot, 2018).

### 2.6 Whole-genome sequence data

We generated low-coverage whole-genome sequence (WGS) data for a subset of 137 *Phyllotis* specimens that were included in the *cytb* data matrix, which we analyzed in conjunction with a chromosome-level reference genome for *Phyllotis vaccarum* (Storz et al., 2023). Depth of coverage ranged from 1.04× to 24.06X (median = 2.58X). According to field identifications and *cytb* haplotypes, this set of specimens represented a total of 11 species (*P. anitae*, *P. camiari*, *P. caprinus*, *P. chilensis*, *P. darwini*, *P. limatus*, *P. magister*, *P. nogalaris*, *P. pehuenche*, *P. vaccarum*, and *P. xanthopygus*), several of which have potentially overlapping ranges (Fig. 1A). All species other than *P. anitae* and *P. nogalaris* are members of the *darwini* species group. Of the 137 vouchered specimens included in the genomic analysis, data for 61 specimens representing *P. chilensis*, *P. limatus*, *P. magister*, and *P. vaccarum* were published previously (Storz et al., 2024).

#### 2.6.1 Genomic library preparation and whole-genome sequencing

All library preparations for whole genome resequencing experiments were conducted in the University using the DNeasy Blood and Tissue kit (Qiagen). We used a Covaris E220 sonicator to shear DNA and we then prepared genomic libraries using the KAPA HyperPlus kit (Roche). Individual libraries were indexed using KAPA UDI’s and pooled libraries were sent to Novogene for Illumina paired-end 150 bp sequencing on a Novaseq X.

#### 2.6.2 Read quality processing and mapping to the reference genome

We used fastp 0.23.2 (Chen et al., 2018) to remove adapter sequences, and to trim and filter low-quality reads from sequences generated from library preparations. We used a 5 bp sliding window to remove bases with a mean quality less than 20 and we discarded all reads <25 bp. We merged all overlapping reads that passed filters and retained all reads that could not be merged or whose paired reads failed filtering. We separately mapped merged reads, unmerged but paired reads, and unpaired reads to the *P. vaccarum* reference genome with BWA 0.7.17 (Li and Durbin, 2009) using the mem algorithm with the -M option which flags split reads as secondary for downstream compatibility. We sorted, merged, and indexed all resulting binary alignment maps with SAMtools 1.15.1 (Li et al., 2009) and used picard 2.27.4 to detect and remove PCR duplicates. We used GATK 3.8 (McKenna et al., 2010) to perform local realignment around targeted indels to generate the final BAM files.

#### 2.6.3 Mitochondrial genome assembly

A *de novo* assembly of the mitochondrial genome of *Phyllotis vaccarum* (specimen UACH8291) as a seed sequence, we used NOVOplasty 4.3.3 (Dierckxsens et al., 2017) to generate *de novo* mitochondrial genome assemblies for all other *Phyllotis* specimens. We annotated assembled mitochondrial genomes with MitoZ to identify coding sequences and we generated a multiple alignment of coding sequence with MAFFT 7.508 (Katoh and Standley, 2013), using the --auto flag to determine the best algorithm given the data.

### 2.7 Analysis of whole-genome sequence variation in *Phyllotis*

First, we randomly downsampled all higher coverage samples to the median coverage (2.58X) using SAMtools 1.17 to avoid artifacts associated with variation in coverage across samples that can impact inferences of population structure. We calculated genotype likelihoods for scaffolds 1-19 (covering >90% of the *Phyllotis* genome) for all samples in ANGSD 0.939 (Korneliussen et al., 2014). We used - GL 2 to specify the GATK model for genotype likelihoods, retained only sites with a probability of being variable >1e-6 with -SNP_pval 1e-6. We filtered out bad and non-uniquely mapped reads with - remove_bads 1 and -uniqueOnly 1, respectively, and only retained reads and bases with a mapping quality higher than 20. We adjusted mapping quality for excessive mismatches with -C 50. We used PCAngsd v.0.99.0 (Meisner and Albrechtsen, 2018) to calculate the covariance matrix from genotype plotted the first, second, and third principal components using the R package *ggplot2* (Wickham, 2016).

Based on results of our genus-wide genomic PCA, we recalculated genotype likelihoods and performed additional genomic analyses on a subset of *P. vaccarum* and *P. limatus* specimens (*n*=51 and 20, respectively). To test for admixture between *P. vaccarum* and *P. limatus*, we calculated ancestry proportions with NGSadmix (Skotte et al., 2013). To alleviate computational costs associated with NGSadmix we generated a reduced SNP set by sampling every hundredth SNP calculated by ANGSD. We ran NGSadmix with K=1-10 with ten iterations for each K value with a random starting seed and a minor allele frequency filter of 0.05. We evaluated the optimal K value using EvalAdmix 0.95 which calculates the pairwise covariance matrix of residuals of model fit. The results of EvalAdmix determined K=2 as the optimal value of K. We combined individual runs for each K value with the R package PopHelper 2.3.1 to average estimates of ancestry across runs.

### 2.8 Genomic patterning of admixture

To examine the genomic patterning of mixed *P. vaccarum*/*P. limatus* ancestry, we conducted a windowed PCA of nucleotide variation. We used the script windowed_pcangsd.py (10.5281/zenodo.8127993) to compute the first principal component in 90% overlapping 1 Mbp windows along chromosomes 1 to 19, using the subset of 51 *P. vaccarum* and 20 *P. limatus* samples and employing minor allele frequency threshold of 0.01. For visualization we excluded outlier windows (those with less than 0.3 % informative sites and those featuring the largest 0.005 % absolute PC1 values across the genome). For consistency we polarized PC1 orientation by its sign for chromosome 1 since polarity is arbitrary in principal component analyses.

## 3. RESULTS

The *cytb* sequence data derive from a total of 448 *Phyllotis* specimens from 169 localities that span most of the distributional range of the genus (Fig. 1B). For the analysis of WGS variation, we used a subset of 137 vouchered specimens representing 11 nominal species of *Phyllotis* that have overlapping or potentially overlapping ranges in Argentina, Bolivia, and Chile. *Phyllotis vaccarum* is one of the most broadly distributed species in this region and different parts of its range potentially overlap with those of *P. caprinus*, *P. chilensis-posticalis*, *P. darwini*, *P. limatus*, *P. magister*, and *P. pehuenche* (Fig. 1A). We therefore concentrated much of our sampling efforts on these zones of range overlap to examine evidence of introgressive hybridization.

### 3.1 Phylogenetic relationships and Divergence Times

At the level of the genus *Phyllotis*, phylogeny estimates based on BI and ML both recovered three Supplementary Figure 1). In the BI analysis, the *andium-amicus* and *osilae* clades were recovered as sister groups (Bayesian Posterior Probability [PP] = 1) (Fig. 2), whereas the ML analysis placed the *osilae* clade as sister to a weaky supported clade (Bootstrap Percentage [BP] = 53) formed by the *andium-amicus* and *darwini* clades (Supplementary Fig. 1). Within the *darwini* group, BI and ML analyses generally recovered the same set of relationships within the *P. xanthopygus* complex, with the exception that the BI phylogeny placed *P. pehuenche* and *P. xanthopygus* as sister (PP = 1; Fig. 2), whereas the ML phylogeny placed *P. xanthopygus* as sister to the clade containing *P. caprinus*, *P. limatus*, *P. vaccarum*, and *P. pehuenche* (BP = 70; Supplementary Fig. 1).

**Figure 2.**
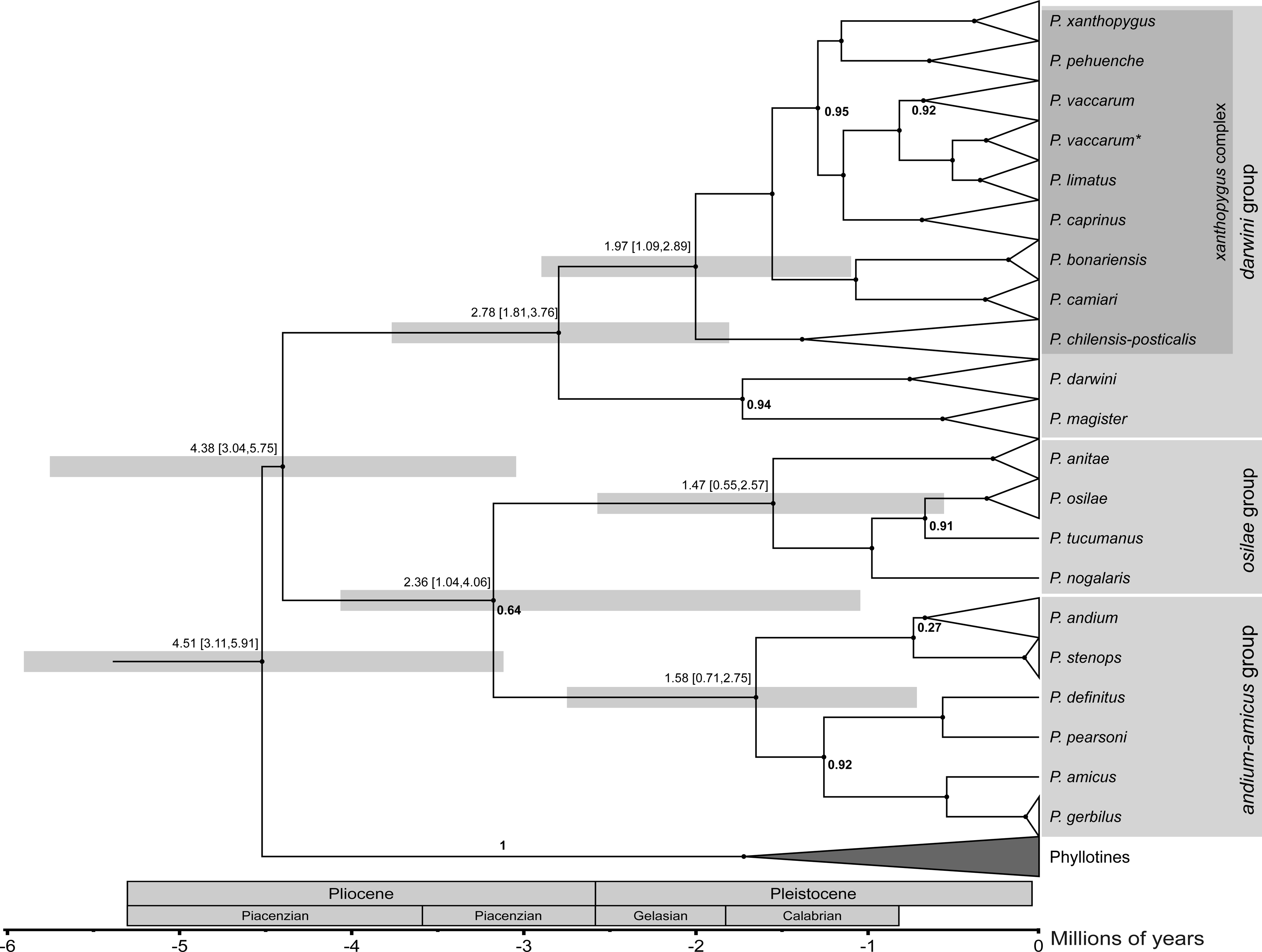
Calibrated maximum clade credibility tree showing Bayesian estimates of phylogenetic relationships and divergence times within the genus *Phyllotis*. Estimates of the 95% Highest Posterior Distributions interval for the divergence times are shown for main clades. Node support is shown only for those cases in which Bayesian posterior probability values were <1. Specimens in the clade labeled ‘*P. vaccarum*\*’ carry *cytb* haplotypes that group with haplotypes of *P. limatus*, even though whole-genome sequence data confirmed their identity as *P. vaccarum* (Storz et al., 2024).

The median estimated crown age for the genus *Phyllotis* was 4.51 Mya with a 95% Highest Posterior Distribution (HPD) of 3.11-5.91 Mya, a range that spans nearly the entire Pliocene. Crown ages and associated HPD’s for the clades corresponding to the species groups *andium-amicus*, *osilae*, and *darwini*, were 1.58 (0.71-2.75), 1.47 (0.55-2.57), and 2.78 Mya (1.81-3.76), respectively. Within each of these three groups, most species diverged during the last ∼2 Mya and there appears to have been a pulse of speciation during the mid to late Pleistocene.

The species delimitation analyses were consistent in recognizing each of the 20 *Phyllotis* taxa represented in the full *cytb* dataset. Different delimitation approaches identified 36-37 distinct units (Fig. 3). Results of the delimitation analyses suggest that *P. caprinus*, *P*. *chilensis*-*posticalis*, *P*. *darwini*, *P*. *magister*, and *P*. *vaccarum* may each represent complexes of multiple species. The internal subdivisions identified within *P*. *caprinus* and *P*. *darwini*, and some of those identified within *P*. *chilensis-posticalis*, have allopatric distributions (Supplementary Fig. 2). Results of the GMYC and PTP delimitation analyses differed in the number of units identified within *P*. *vaccarum* and *P*. *pehuenche*. The GMYC and b-GMYC analyses identified six distinct units within *P*. *vaccarum* and recognized *P*. *pehuenche* as a single unit. By contrast, the PTP and b-PTP approaches recognized three distinct units within both *P*. *vaccarum* and *P*. *pehuenche*.

**Figure 3.**
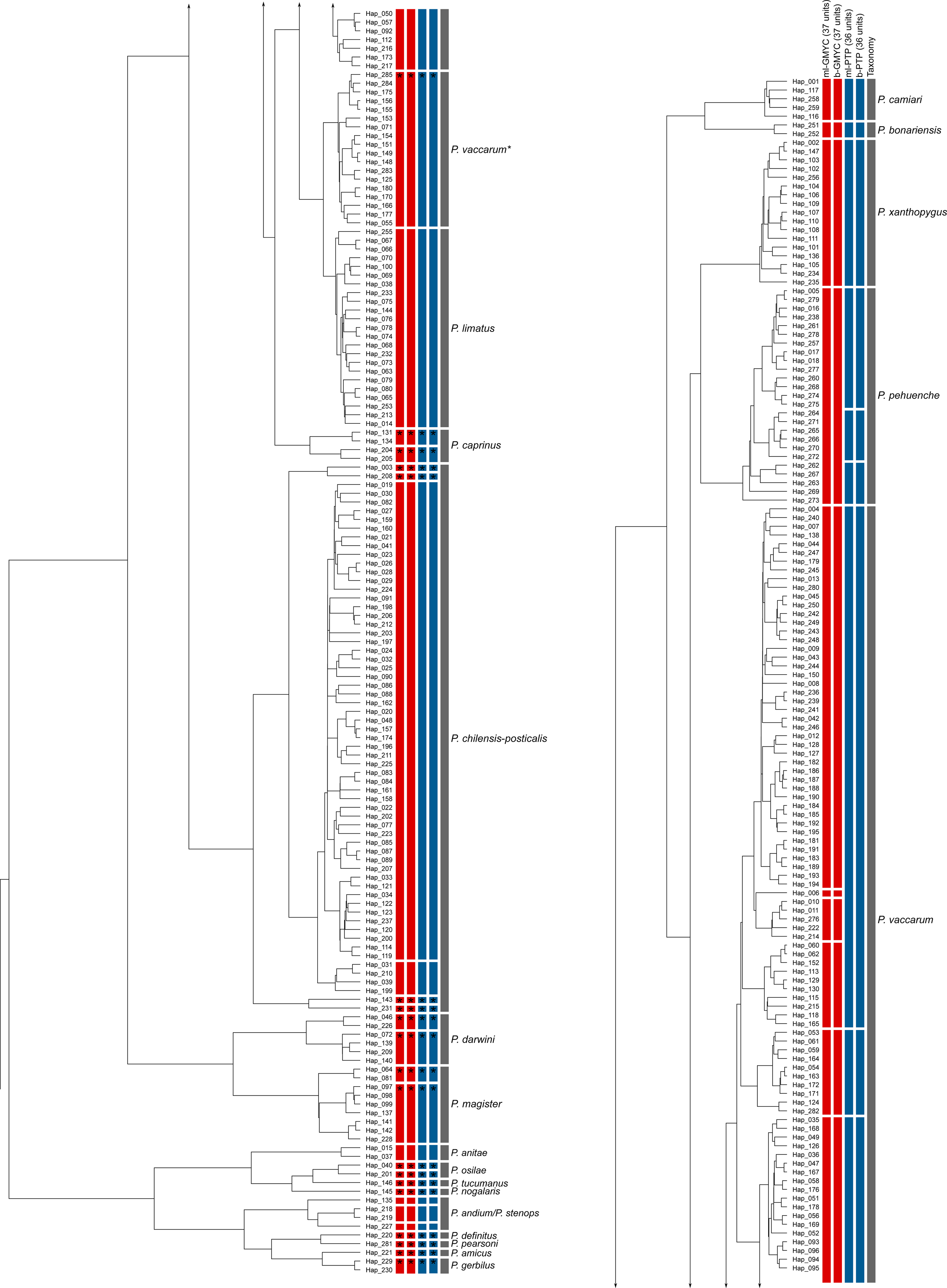
Maximum clade credibility depicting the delimitation schemes inferred from GMYC (red bars) and PTP (blue bars). Gaps in the vertical bars denote units delimited by each method, and asterisks denote splits with support values >0.75. Continuous gray bars denote current taxonomic designations for nominal species. Terminal labels depict the haplotype classes of sequences that were retained to construct the non-redundant matrix of *cytb* haplotypes. Specimens in the clade labeled ‘*P. vaccarum*\*’ carry *cytb* haplotypes that group with haplotypes of *P. limatus*, even though whole-genome sequence data confirmed their identity as *P. vaccarum* (Storz et al., 2024).

Levels of cytb sequence differentiation between pairs of *Phyllotis* species are highly variable, with estimated *p*-distances ranging from 2.73% (SE = 0.004) between *P*. *limatus* and *P*. *vaccarum*, to 17.28% (SE = 0.013) between *P*. *gerbilus* and *P*. *nogalaris* (Table 1). The mean *p*-distance between nominal species within the genus *Phyllotis* is 7.55% (SE = 0.005). Within the *Phyllotis xanthopygus* species complex, the maximum *p*-distance is 10.82% between *P*. *pehuenche* and *P*. *chilensis* (Table 1). We also estimated *p*-distances between internal subdivisions (candidate species) within several nominal forms that were identified as significant in the species delimitation analyses. In these cases, pairwise *p*-distances ranged from 1.81% (SE = 0.003) between subdivisions within *P*. *magister* to 9.32% (SE = 0.011) between the most divergent subdivisions within *P*. *chilensis* (Supplementary Table 2).

**Table 1.** Mean *cytb p*-distances between pair of species of *Phyllotis* (below diagonal). Mean values for intraspecific *p*-distances are shown in bold on the diagonal. Standard errors (SE) for each estimate of pairwise distance is shown above the diagonal.

|  | 1 | 2 | 3 | 4 | 5 | 6 | 7 | 8 | 9 | 10 | 11 | 12 | 13 | 14 | 15 | 16 | 17 | 18 | 19 | 20 |
| --- | --- | --- | --- | --- | --- | --- | --- | --- | --- | --- | --- | --- | --- | --- | --- | --- | --- | --- | --- | --- |
| 1. *P. amicus* | -- | 0.998 | 1.221 | 1.205 | 1.204 | 1.178 | 1.241 | 1.219 | 1.011 | 1.114 | 1.255 | 1.289 | 1.223 | 1.154 | 1.165 | 1.050 | 1.121 | 1.215 | 1.159 | 1.303 |
| 2. *P. andium* | 11.498 | **5.381** | 1.193 | 1.193 | 1.195 | 1.082 | 1.075 | 1.061 | 1.111 | 1.125 | 1.049 | 1.260 | 1.138 | 0.920 | 1.234 | 1.112 | 0.559 | 1.286 | 1.120 | 1.134 |
| 3. *P. anitae* | 14.414 | 12.032 | **1.253** | 1.335 | 1.237 | 1.355 | 1.166 | 1.347 | 1.398 | 1.387 | 1.259 | 1.083 | 1.078 | 1.156 | 1.482 | 1.322 | 1.224 | 1.071 | 1.366 | 1.343 |
| 4. *P. bonariensis* | 15.855 | 13.615 | 14.052 | **0.749** | 0.877 | 0.864 | 1.044 | 1.238 | 1.180 | 0.978 | 0.964 | 1.276 | 1.252 | 1.132 | 0.879 | 1.022 | 1.140 | 1.315 | 0.886 | 0.880 |
| 5. *P. camiari* | 14.657 | 13.048 | 12.119 | 8.514 | **1.049** | 0.886 | 1.000 | 1.244 | 1.256 | 1.036 | 0.986 | 1.214 | 1.225 | 1.160 | 1.001 | 1.019 | 1.306 | 1.290 | 0.988 | 0.938 |
| 6. *P. caprinus* | 16.062 | 13.625 | 13.616 | 8.704 | 9.218 | **4.061** | 1.072 | 1.207 | 1.260 | 0.822 | 1.034 | 1.288 | 1.178 | 1.146 | 0.911 | 0.877 | 1.215 | 1.148 | 0.787 | 0.914 |
| 7. *P. darwini* | 15.844 | 13.466 | 13.597 | 12.196 | 12.774 | 12.469 | **3.305** | 1.334 | 1.318 | 1.020 | 1.062 | 1.254 | 1.198 | 1.170 | 1.116 | 1.133 | 1.235 | 1.178 | 1.016 | 1.079 |
| 8. *P. definitus* | 12.453 | 10.873 | 14.994 | 15.105 | 15.341 | 15.807 | 15.459 | **0.001** | 1.280 | 1.269 | 1.460 | 1.347 | 1.391 | 0.995 | 1.354 | 1.385 | 1.125 | 1.327 | 1.232 | 1.234 |
| 9. *P. gerbilus* | 6.173 | 12.031 | 15.722 | 15.432 | 15.240 | 15.931 | 16.540 | 12.638 | **0.274** | 1.211 | 1.302 | 1.486 | 1.385 | 1.236 | 1.135 | 1.174 | 1.269 | 1.270 | 1.219 | 1.337 |
| 10. *P. limatus* | 14.237 | 12.813 | 13.227 | 8.636 | 9.014 | 7.250 | 11.946 | 14.827 | 14.435 | **0.512** | 1.021 | 1.350 | 1.324 | 1.052 | 0.976 | 0.999 | 1.284 | 1.283 | 0.396 | 1.046 |
| 11. *P. magister* | 14.915 | 12.630 | 13.351 | 10.609 | 10.575 | 11.008 | 10.852 | 14.961 | 15.877 | 9.711 | **1.568** | 1.139 | 1.082 | 1.176 | 1.030 | 1.002 | 1.125 | 1.135 | 0.976 | 0.956 |
| 12. *P. nogalaris* | 16.105 | 14.206 | 11.259 | 14.232 | 14.157 | 16.030 | 15.114 | 15.698 | 17.284 | 14.566 | 14.328 | -- | 1.035 | 1.146 | 1.373 | 1.352 | 1.302 | 1.067 | 1.296 | 1.320 |
| 13. *P. osilae* | 14.723 | 12.551 | 10.068 | 14.082 | 13.900 | 14.708 | 15.349 | 15.721 | 16.283 | 14.043 | 13.301 | 10.205 | **3.125** | 1.158 | 1.331 | 1.252 | 1.185 | 0.910 | 1.309 | 1.197 |
| 14. *P. pearsoni* | 12.406 | 9.862 | 13.181 | 14.286 | 14.361 | 15.353 | 14.387 | 7.103 | 12.948 | 13.878 | 13.234 | 15.664 | 14.210 | -- | 1.214 | 1.118 | 0.952 | 1.202 | 0.986 | 1.158 |
| 15. *P. pehuenche* | 15.874 | 14.086 | 14.730 | 9.277 | 10.602 | 9.128 | 13.325 | 16.252 | 16.280 | 8.894 | 11.256 | 15.689 | 15.516 | 15.499 | **1.449** | 0.950 | 1.270 | 1.279 | 0.961 | 1.009 |
| 16. *P. chilensis-posticalis* | 15.236 | 14.469 | 14.023 | 9.507 | 9.632 | 9.674 | 12.576 | 16.412 | 15.647 | 9.095 | 10.992 | 14.492 | 15.134 | 14.687 | 10.820 | **1.578** | 1.180 | 1.172 | 0.980 | 0.997 |
| 17. *P. stenops* | 11.857 | 4.801 | 11.710 | 13.111 | 12.797 | 13.393 | 13.883 | 11.093 | 12.250 | 12.668 | 11.744 | 14.680 | 11.732 | 10.056 | 14.407 | 14.184 | **0.252** | 1.289 | 1.266 | 1.246 |
| 18. *P. tucumanus* | 14.591 | 12.660 | 10.032 | 13.836 | 13.182 | 14.074 | 14.370 | 14.571 | 15.768 | 14.041 | 12.413 | 10.189 | 6.811 | 13.962 | 15.467 | 14.241 | 11.909 | -- | 1.328 | 1.338 |
| 19. *P. vaccarum* | 15.351 | 13.194 | 14.289 | 8.512 | 9.186 | 7.304 | 12.260 | 15.270 | 15.453 | 2.733 | 10.171 | 14.894 | 14.756 | 13.973 | 9.170 | 9.513 | 13.164 | 14.946 | **2.224** | 0.981 |
| 20. *P. xanthopygus* | 15.205 | 12.941 | 14.237 | 8.010 | 9.383 | 8.304 | 11.199 | 15.837 | 15.032 | 8.668 | 10.461 | 14.008 | 13.467 | 14.902 | 9.540 | 10.279 | 12.910 | 13.890 | 8.737 | **0.829** |

### 3.2 Genomic assessment of species limits

To further examine species limits suggested by the analysis of *cytb* sequence variation, we generated low-coverage WGS data for representative subsets of specimens from 11 nominal species, several of which have overlapping ranges in the Altiplano and/or adjoining lowlands. We also derived an alignment of whole mitochondrial genomes from the WGS data. Whereas the BI and ML analyses of *cytb* variation yielded some conflicting estimates of species relationships within the *P. xanthopygus* species complex (Fig. 2 and Supplementary Fig. 1), the ML phylogeny estimate based on complete mitochondrial genomes found that *P. pehuenche* and *P. xanthopygus* are sister to each other and placed them sister to the clade comprising *P. caprinus*, *P. limatus*, and *P. vaccarum* (BP = 100) (Fig. 4).

**Figure 4.**
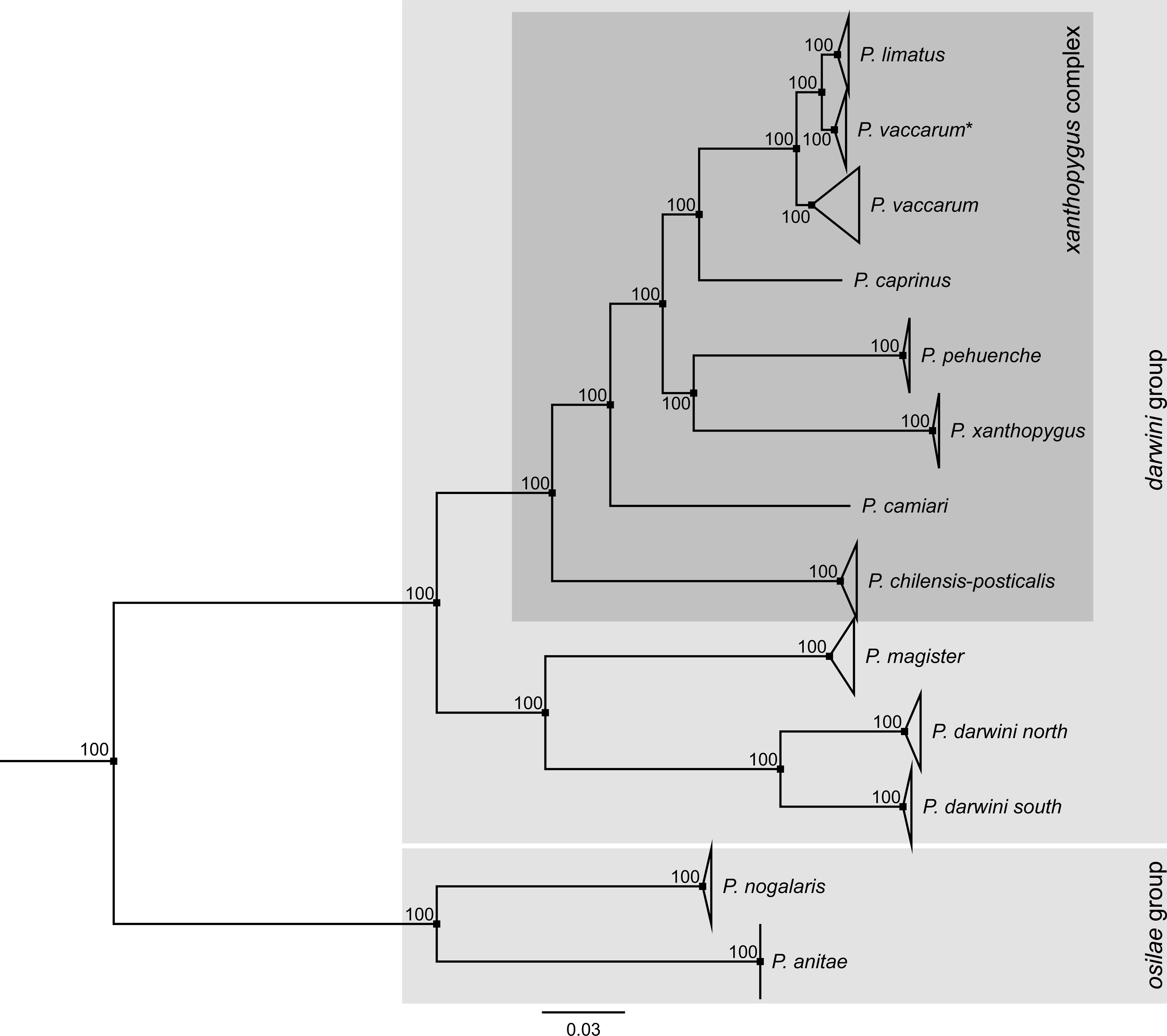
Maximum likelihood tree estimated from coding sequence of complete mitochondrial genomes for a set of 11 nominal *Phyllotis* species. Numbers adjacent to internal nodes denote ultrafast bootstrap support values for each clade. Within the taxon currently recognized as *P. darwini*, the species delimitation analysis identified two highly distinct subdivisions (see Fig. 3). Representatives of both internal subdivisions form distinct clades in the mitogenome tree, which we labeled ‘*P. darwini* south’ and ‘*P. darwini*’ north.

In a PCA of genome-wide variation, PC1, PC2, and PC3 captured 36.8%. 23.2%, and 7.15% of the total variation, respectively (Fig. 5A,B). Samples of *P. darwini* from the northern and southern portions of the species range separated into two highly distinct clusters (Fig. 5A,B). The distinct clusters of *P*. *darwini* specimens identified in the genomic PCA are fully congruent with two divergent mtDNA clades that were identified as significant internal subdivisions in the species delimitation analysis (Figure 3). On the basis of *cytb* sequence data, the estimated *p*-distance between the northern and southern subdivisions of *P. darwini* was 5.40% (SE = 0.679) (Supplementary Table 2).

**Figure 5.**
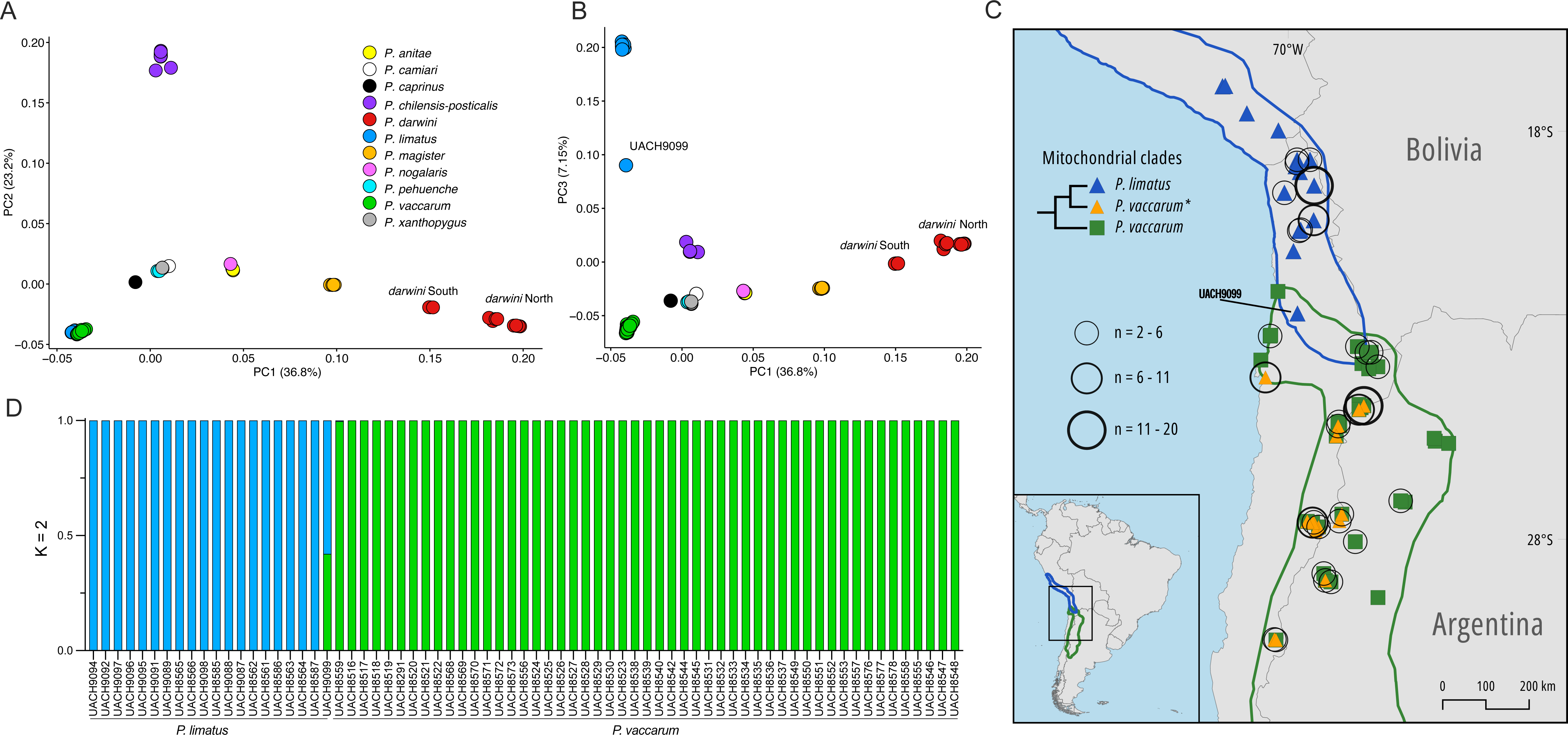
Genomic variation among species of *Phyllotis* based on 137 samples representing 11 nominal species. A) Genomic principal component analysis (PCA) of genome-wide variation (PC1 vs PC2). Two distinct clusters of nominal *P. darwini* specimens, ‘*darwini* South’ and ‘*darwini* North’, are distinguished along the PC1 axis. B) Plot of PC1 vs PC3 separates *P. limatus* and *P. vaccarum* along the PC3 axis, and reveals a single specimen, UACH9099 (designated *P. limatus* based on mtDNA haplotype), which has a PC3 score intermediate between the two species. C) Map of collecting localities and distribution limits of *P. limatus* and *P. vaccarum*. UACH9099 comes from a site located in a narrow zone of range overlap between the two species in northern Chile. The map also shows the distribution of mice that are identified as *P. vaccarum* on the basis of whole-genome sequence data, but which carry mtDNA haplotypes that are more closely related to those of *P. limatus* (denoted as ‘*P. vaccarum*\*’ in the inset tree diagram). D) Structure plot showing clear distinction between *P. limatus* and *P. vaccarum* (*n*=20 and 51, respectively). The putative hybrid specimen, UACH9099, was assigned almost exactly equal ancestry proportions from the two species.

Using coding sequence of the complete mitochondrial genome, the corresponding *p*-distance was 7.25% (SE = 0.002).

The sister species *P. limatus* and *P. vaccarum* were not readily distinguishable along the first two PC axes (Fig. 5A), but they were cleanly separated along PC3 (Fig. 5B). One specimen, UACH9099, which was identified as *P. limatus* on the basis of mtDNA, fell in between the two distinct clusters of *P. limatus* and *P. vaccarum* samples along PC3 (Fig. 5B). Specimen UACH9099 was collected in the narrow zone of range overlap between southern *P. limatus* and northern *P. vaccarum*, about 200-250 km from the localities where *P. vaccarum* specimens were found to carry mtDNA haplotypes more closely related to those of *P. limatus* than to other *vaccarum* (Fig. 5C). Individual admixture proportions estimated with NGSadmix also distinguished *P. limatus* and *P. vaccarum* samples as genetically distinct clusters, and UACH9099 was assigned approximately equal admixture proportions of the two species (Fig. 5D). A sliding window analysis of PC1 comprising the full sample of *P. limatus* and *P. vaccarum* specimens revealed a mosaic patterning of variation along the genome of UACH9099, as autosomal segments alternated between three main patterns: (*i*) homozygous for *P. limatus* ancestry, (*ii*) homozygous for *P. vaccarum* ancestry, or (*iii*) heterozygous, falling approximately halfway in between the two species (Fig. 6).

**Figure 6.**
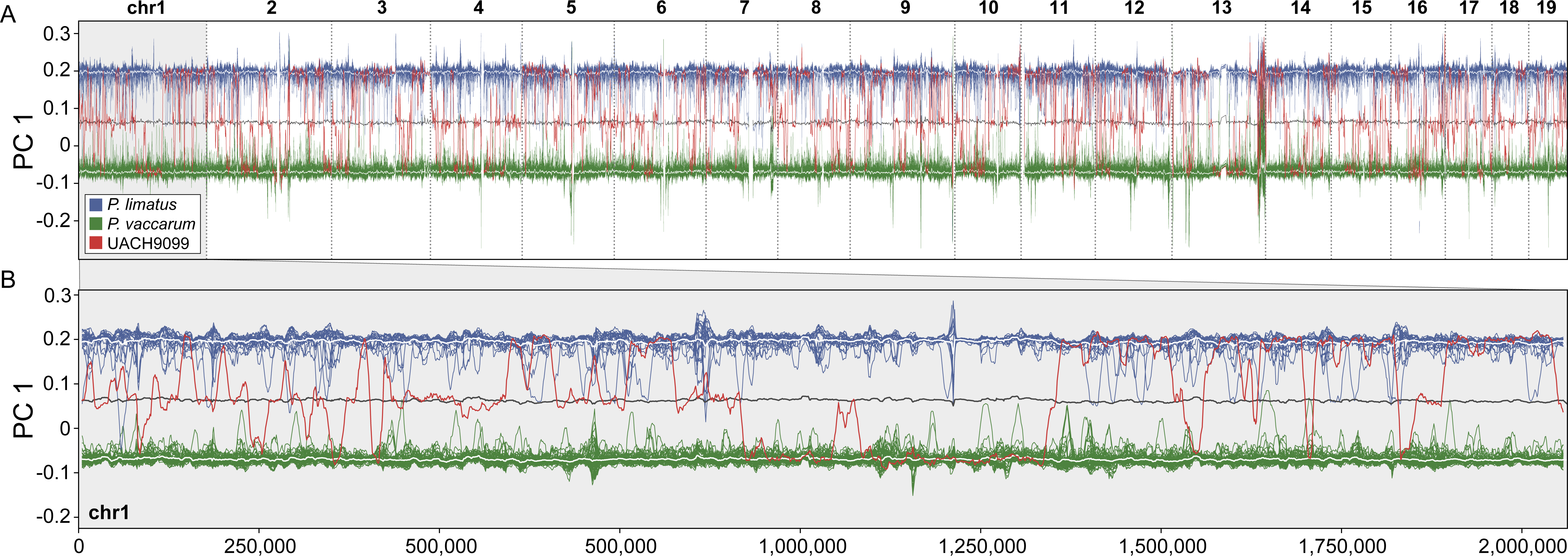
Windowed PCA of a *P. vaccarum* x *P. limatus* hybrid. PC1 was computed in overlapping 1 Mbp windows along the genome for a subset of 50 *P. vaccarum* (green), 20 *P. limatus* (blue), and the putative hybrid, UACH9099 (red). Mean PC1 values for each species are shown as white lines and the mean value between both species’ averages is shown as a grey line. UACH9099 features a mosaic genome, with its local ancestry alternating between *P. vaccarum, P. limatus*, or a point intermediate between the two species. (A) Windowed PCA of chromosomes 1-19. (B) High resolution visualization of PC 1 along chromosome 1.

### 3.3 Revised geographic Range Limits of *Phyllotis* Species

The integrated analysis of mtDNA and WGS data enabled us to delineate the geographic range limits *caprinus* that we collected in southern Bolivia significantly extend the species’ known range to the north (Fig. 7). Another possibility suggested by results of the species delimitation analysis (Fig. 3) is that the specimens from central Bolivian do not represent extralimital records of *P. caprinus*, but may instead represent a new, undescribed species sister to the form *P. caprinus* that distributes in southern Bolivia and northern Argentina. In the case of *P*. *chilensis-posticalis*, our specimens from the Chilean regions of Arica y Parinacota, Tarapacá, and Antofagasta extend the species’ known range to the west (Fig. 7).

**Figure 7.**
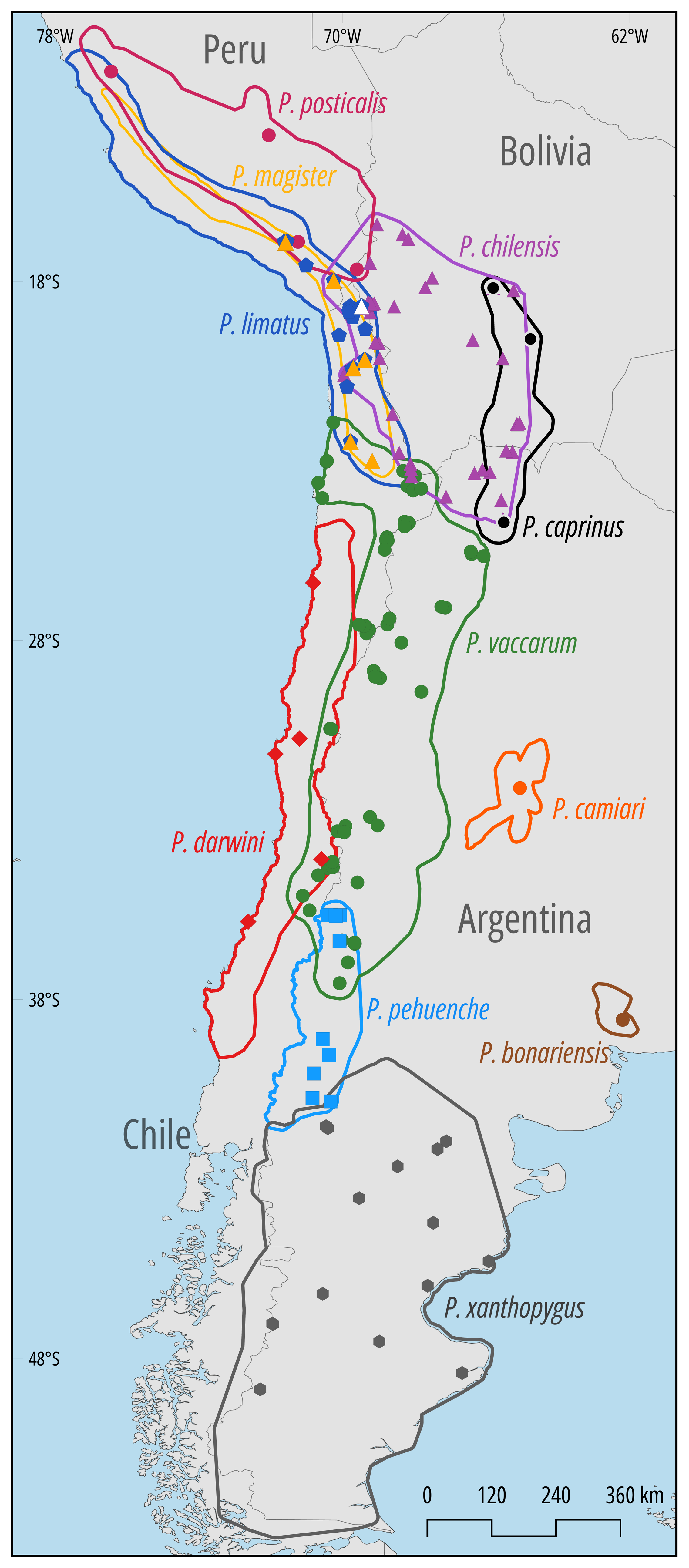
Revised distribution limits of species in the *Phyllotis darwini* species group based on mtDNA and WGS data. Filled circles denote collection localities that helped define geographic range limits.

Our records for *P*. *vaccarum* indicate that this primarily highland species is replaced by *P. darwini* at elevations <2000 m on the western slope of the Andes, but – beyond the northernmost limits of *P. darwini* – the range of *P*. *vaccarum* extends all the way to sea level along a narrow stretch of coastline in northern Chile (Fig. 7). On the eastern slope of the Andes, our records from northwestern Argentina indicate that the species does not occur below 1200 m, as it is replaced by *P. anitae* and *P. nogalaris* in lowland Yungas forests. Further south along the eastern slope of the Cordillera where humid lowland forests give way to arid steppe and Monte habitats, our lowest elevation records of *P. vaccarum* were from 765-1158 m in the Argentine provinces of Catamarca, Neuquén, and Mendoza, but the majority of records are from elevations above 1200 m.

## 4. DISCUSSION

### 4.1 Most diversification of *Phyllotis* occurred in the Pleistocene

Estimating divergence times of Sigmodontine rodents has been difficult due to a lack of suitable fossils that could be used to calibrate molecular data (Salazar-Bravo et al., 2013). Previous studies, using a maximum likelihood clock estimate of 7.3% divergence per Mya (Steppan et al., 2004, 2007), placed the basal split of *Phyllotis* in the Pliocene (3.0–5.1 Mya) and the basal split of the *P. xanthopygus* species complex in the Pliocene-Pleistocene transition (1.6–2.3 Mya). Riverón (2011) estimated a similar Pliocene basal split for *Phyllotis* (2.83-4.05 Mya) using an analogous strict-clock estimate. Our secondary calibration-based estimations suggest a similar timing of diversification of *Phyllotis*, with an estimated initial divergence 4.51 Mya (95% HPD = 3.11-5.91 Mya) and subsequent diversification of the *P. xanthopygus* complex 2.78 Mya (95% HPD = 1.81-3.76 Mya). However, divergence time estimates should be always interpreted with caution due to uncertainty about the calibration approaches employed and the taxon sampling used in the phylogeny estimate (Steppan et al., 2007; Parham et al., 2012).

In principle, the diversification of *Phyllotis* could have been spurred by mountain uplift and/or climate-related environmental changes at the end of the Pliocene and the beginning of the Pleistocene. The Central Andean Plateau experienced the most significant phase of uplift in the late diversification of *Phyllotis* would have started well before the end of the Pliocene (2.6 Mya). It is also possible that diversification occurred more recently, and independently of Andean uplift, during periods of climate-induced environmental change in the Pleistocene. For example, the mid-Pleistocene Transition (MPT; 1.25–0.70 Mya) was associated with a major shift in global climate periodicity that produced a persistent global aridification trend (Herbert, 2023). Thus, the Pleistocene Aridification hypothesis predicts that diversification of *Phyllotis* would have occurred more recently than the Andean uplift, coinciding with periods of climate change that were not directly related to orogenic events.

Our results suggest that most diversification of *Phyllotis* occurred during the past 3 million years with divergence times for most species coinciding with glacial cycles in the mid- to late Pleistocene (Fig. 2). Basal splits in two of the three main *Phyllotis* clades (the *andium-amicus* and *osilae* species groups) occurred prior to the MPT (0.7-1.25 Mya), whereas the basal split within the *darwini* group is estimated to have occurred 2.78 Mya (95% HPD = 1.81-3.76 Mya) close to the Pliocene-Pleistocene boundary. Within each of the three main clades, most diversification occurred within the past ∼1.47-1.97 Mya. Thus, our results suggest that most diversification of *Phyllotis* occurred well after the late Miocene-Pliocene phase of Andean uplift.

### 4.2 Alpha diversity within the *Phyllotis darwini* species group

Based on results of our phylogenetic reconstructions and species delimitation analyses, we can identify at least 10 lineages that are referable to traditionally recognized forms within the *Phyllotis darwini* species group (Figs. 2, 3, and 4). However, results of the species delimitation analysis clearly show that some of these nominal forms may encompass more than one species. There appears to be potential for the existence of additional species within nominal forms that are currently recognized as *P*. *caprinus*, *P*. *chilensis-posticalis*, *P*. *darwini*, *P*. *magister*, and *P*. *vaccarum* (Fig. 3). The distinction of these candidate species requires further taxonomic work.

The Bolivian specimens of *P*. *caprinus* from Chuquisaca (MSB237236) and Cochabamba (MSB238568) constitute a clade with a high degree of differentiation relative to the remaining Argentine specimens that are referable to typical *P. caprinus* (*cytb p*-distance=5.6%, SE=0.008) (Figure 3). *Phyllotis darwini* and *P*. *chilensis-posticalis* also exhibit north-south patterns of internal structure (Supplementary Fig. S2A,B), with highly distinct units identified by the species delimitation analyses (Fig. 3). In the case of *P. darwini*, divergence between northern and southern mtDNA clades is also apparent at the whole-genome level (Fig. 5A,B). Consistent with results of Ojeda et al. (2021), the clade that includes specimens that we refer to as *P. chilensis-posticalis* appears likely to contain multiple cryptic species with apparently allopatric distributions in Peru (Supplementary Fig. 2B).

Although Ojeda et al. (2021) referred to this group as the “*P*. *posticalis-rupestris*” clade, geographic with the southern-most distribution in northeastern Chile, southwestern Bolivia, and northwestern Argentina (Hershkovitz, 1962; Mann, 1945; Thomas, 1912; Supplementary Fig. S2B). Here and elsewhere (Storz et al., 2014), we followed Mann (1945) and Pearson (1958) in using the name *P*. *chilensis* for the mice in this subclade that we collected in the Altiplano of northern Chile, southwestern Boliva, and northwestern Argentina. Therefore, we prioritize the use of *P. posticalis* for the subclade with the northern-most distribution as it includes a specimen from the vicinity of the associated type locality in the Department of Junín, Peru (Thomas, 1912). The distinction of these lineages (Fig. 3) requires further analysis using morphological and genomic data.

In *P. vaccarum*, one *cytb* haplogroup that was identified as a distinct unit in the species delimitation analysis is sister to a clade formed by haplotypes of *P. limatus*. The *P. vaccarum* mice that harbor *limatus*-like mtDNA haplotypes are not distinguishable from other *P. vaccarum* at the whole-genome level (Storz et al. 2024). In this particular case of mitonuclear discordance, identified mtDNA subdivisions are clearly not reflective of cryptic species within *P. vaccarum*.

### 4.3 Evidence for interspecific hybridization

The genomic data revealed clear-cut evidence of ongoing hybridization between *P. limatus* and *P. vaccarum* (Fig. 5D and Fig. 6), suggesting that introgression is a plausible explanation for the observed mitonuclear discordance between the two species (Fig. 5C; see Storz et al., 2024). The UACH9099 specimen carries *P. limatus* mtDNA but harbors approximately equal genome-wide admixture proportions from *P. limatus* and *P. vaccarum* (Fig. 5D). At face value, the approximately equal admixture proportions suggest that UACH9099 could be a first generation (F1) interspecific hybrid that has received one haploid complement of chromosomes from each parental species.

However, in the windowed PCA, an F1 hybrid would be expected to localize halfway between the two divergent parental stocks. Contrary to that expectation, tracts across the genome of UACH9099 were either homozygous for *P. vaccarum* ancestry, homozygous for *P. limatus* ancestry, or heterozygous (i.e., combining both species’ genomes) (Fig. 6). The mosaic patterning of nucleotide variation appears to reflect one or more rounds of recombination subsequent to an initial *P. limatus* x *P. vaccarum* hybridization event and suggests that UACH9099 is the product of an F2 or more advanced-stage intercross. Given that UACH9099 was assigned roughly equal admixture proportions for both species (Fig. 5C), it is likely that the zone of range overlap between *P. limatus* and *P. vaccarum* in northern Chile represents a zone of ongoing hybridization. Although the observed pattern of genomic mosaicism in UACH9099 could have been produced by a balanced number of backcrossing events with both parental species, we regard ongoing mattings between hybrids as a more likely scenario. More intensive collecting from the zone of range overlap between *P. limatus* and *P. vaccarum* will be required to assess the pervasiveness of hybridization between the two species. *P. vaccarum*, which also happen to be the only pair of sister species with overlapping ranges within the *P. darwini* group, all remaining *Phyllotis* specimens that grouped together in the *cytb* phylogeny were also identified as distinct groupings in the analysis of WGS data (Fig. 5A,B).

### 4.4 A revised understanding of geographical range limits of *Phyllotis* mice

The use of sequence data to confirm the identities of all collected specimens provided new information about geographic range limits and revealed notable range extensions for several species of *Phyllotis* (Fig. 7). The westward range extension of *P. chilensis* in northern Chile is noteworthy because only *P*. *limatus* and *P*. *magister* had been previously recorded in this zone (Steppan and Ramírez, 2015: Ojeda et al., 2021). We collected specimens referable to *P. chilensis* from several extremely high- elevation localities in northern Chile and western Bolivia, including multiple specimens from 5221 m on the flanks of Volcán Parinacota and 5027 m on the flanks of Volcán Acotango in western Bolivia. Such records highlight the importance of surveying environmental extremes to accurately characterize geographic range limits, especially for taxa like *Phyllotis* that are known to inhabit extreme southern latitudes in Patagonia, extreme elevations in the Central Andes, and extreme arid zones in the Atacama Desert. *P. vaccarum* was previously documented to have the broadest elevational range of any mammal, from the coastal desert of northern Chile to the summits of >6700 m volcanoes (Storz et al., 2020, 2024). The species has a similarly broad elevational range on the eastern slope of the Andes, but the lower range limit depends on the nature of the low elevation biome (Jayat et al., 2021; Riverón, 2011). In northwest Argentina, the species appears to have a lower range limit >1200 m, as it is replaced by species in the *osilae* group in humid Yungas forests. In central western Argentina, *P. vaccarum* reaches elevations below 1000 m in arid Patagonian steppe and Monte habitats.

## 5. Conclusions

Our intensive collecting in the Andean Altiplano and surrounding lowlands enabled us to fill key gaps in geographic coverage. By integrating vouchered specimen records with species identifications based on phenotypic, mtDNA and WGS data, we now have a better understanding of geographic range limits for species of the *P. darwini* group. The delimitation of genetically distinct units within several named forms indicates the presence of much undescribed alpha diversity in *Phyllotis*, as pointed out by previous authors (e.g., Ojeda et al., 2021; Jayat et al., 2021). Within the *P. xanthopygus* complex, *P. limatus* and *P. vaccarum* represent the only species for which we observed mitonuclear discordance and documented ongoing hybridization. This example indicates that interspecific hybridization occurs in *Phyllotis*, but more intensive collecting in zones of range overlap between species will be required to assess the pervasiveness of introgressive hybridization in the group.

Although much of the diversification of *Phyllotis* may have occurred in the Andean highlands, our phase of uplift of the Central Andean Plateau in the Miocene-late Pliocene. Instead, most lineage splitting seems to be associated with climatically induced environmental changes in the mid- to late Pleistocene.

## AUTHOR CONTRIBUTIONS

MQ-C, GD, and JFS designed the study, MQ-C, NMB, GD, PJ, PT, and JFS performed the fieldwork, SL, JLM, and TM performed the laboratory work, MQ-C, SL, JLM, TM, JAC, LMB, NDH, ZAC, JMG, GD, and JFS performed data analysis and/or helped with interpretation, MQ-C and JFS wrote the initial draft of the manuscript, and all authors read and approved it.

## ACKNOWLEDGMENTS

We thank Mario Pérez-Mamani and Juan Carlos Briceño for assistance and companionship in the field, Alex González for assistance in the lab, and José Urquizo for helpful comments and discussion.

## DATA AVAILABILITY STATEMENT

The genomic data associated with this study are openly available in the NCBI bioproject PRJNA950396. The newly generated *cytb* sequences are available in GenBank (accession numbers: PQ295377-PQ295555)

## ETHICS STATEMENT

All animals were collected in the field with permission from the following agencies: Servicio Agrícola y Ganadero, Chile (6633/2020, 2373/2021, 5799/2021, 3204/2022, 3565/2022, 911/2023 and 7736/2023), Corporación Nacional Forestal, Chile (171219, 1501221, and 31362839), Dirección Nacional de Fronteras y Límites del Estado, Chile (DIFROL, Autorización de Expedición Científica #68 and 02/22), Ministerio de Medio Ambiente y Agua, Estado Plurinacional de Bolivia (Resolución Administrativa 026/09 and DVS-CRT-02/91), and the Secretaria de Ambiente (Ministerio de Desarrollo Sustentable) de Jujuy, and the Ministerio de Desarrollo Productivo (Direccion de Flora, Fauna Silvestre y Suelos) de Tucumán, Argentina (Expte. N° P4-00402-21 Disp. S.A. N° 001/22, Expte. N° P4 -00158 -22 Disp. S.A. N° 007/22, Expte. N° 677-330-2021, and Expte. N° 677-330-2021). All live-trapped animals were handled in accordance with protocols approved by the Institutional Animal Care and Use Committee (IACUC) of the University of Nebraska (project ID’s: 1919, 2100), IACUC of the University of New Mexico (project ID’s: 16787 and 20405), and the bioethics committee of the Universidad Austral de Chile (certificate 456/2022).

