## Supplementary Material for "Species limits and hybridization in Andean leaf-eared mice (*Phyllotis*)"

Table S1

Figures S1 and S2

**Table S1**. Mean *p*-distance between pairs of lineages within nominal species of *Phyllotis* that were identified by the species delimitation analyses. Mean intra-lineage distance estimates are shown in bold in the diagonal. Standard errors (SE) of each pairwise distance estimate is shown above the diagonal. Internal clades are named according to their position (top-to-bottom) along branch-tips of the tree depicted in Figure 2. Specimens in the group labeled '*P. vaccarum**' carry *cytb* haplotypes that group with haplotypes of *P. limatus*, even though whole-genome sequence data confirmed their identity as *P. vaccarum* (Storz et al., 2024).

| | 1 | 2 | 3 | 4 | 5 | 6 | 7 | 8 | 9 | 10 | 11 | 12 | 13 | 14 |
|---|---|---|---|---|---|---|---|---|---|---|---|---|---|---|
| 1. *P. caprinus* 1 | **0.379** | 0.770 | 1.047 | 1.208 | 1.232 | 1.139 | 1.094 | 1.101 | 0.945 | 0.911 | 1.016 | 1.086 | 0.901 | 0.927 |
| 2. *P. caprinus* 2 | 5.591 | **1.623** | 1.133 | 1.265 | 1.064 | 1.046 | 1.116 | 1.049 | 1.014 | 1.037 | 1.122 | 1.152 | 0.831 | 0.886 |
| 3. *P. darwini* 1 | 11.976 | 11.549 | **4.061** | 0.679 | 1.173 | 1.178 | 1.114 | 1.152 | 1.115 | 1.062 | 1.145 | 1.253 | 0.979 | 1.048 |
| 4. *P. darwini* 2 | 13.285 | 12.360 | 5.399 | **0.915** | 1.128 | 1.100 | 1.201 | 1.224 | 1.239 | 1.230 | 1.133 | 1.288 | 1.110 | 1.127 |
| 5. *P. magister* 1 | 11.772 | 10.637 | 10.508 | 10.762 | **0.166** | 0.348 | 0.987 | 0.960 | 1.064 | 1.070 | 1.156 | 1.069 | 0.999 | 1.010 |
| 6. *P. magister* 2 | 11.277 | 10.620 | 10.924 | 10.895 | 1.809 | **1.507** | 0.940 | 0.869 | 1.023 | 1.041 | 1.053 | 1.005 | 0.996 | 1.004 |
| 7. *P. chilensis/posticalis* 1 | 10.113 | 10.674 | 12.616 | 12.890 | 10.824 | 10.470 | **--** | 0.661 | 0.790 | 0.820 | 0.953 | 0.943 | 1.141 | 1.153 |
| 8. *P. chilensis/posticalis* 2 | 9.987 | 9.925 | 11.863 | 12.079 | 8.951 | 8.483 | 3.995 | **--** | 0.680 | 0.637 | 0.986 | 0.978 | 1.049 | 1.010 |
| 9. *P. chilensis/posticalis* 3 | 9.475 | 9.799 | 12.294 | 12.717 | 11.321 | 10.934 | 6.122 | 5.116 | **0.888** | 0.386 | 0.981 | 1.036 | 1.003 | 1.021 |
| 10. *P. chilensis/posticalis* 4 | 9.438 | 9.785 | 12.397 | 12.633 | 11.511 | 11.126 | 6.367 | 5.056 | 2.069 | **1.132** | 0.993 | 1.054 | 1.022 | 1.057 |
| 11. *P. chilensis/posticalis* 5 | 10.407 | 10.025 | 12.491 | 12.421 | 10.051 | 9.919 | 8.701 | 7.818 | 8.994 | 9.001 | **--** | 0.966 | 1.215 | 1.228 |
| 12. *P. chilensis/posticalis* 6 | 11.809 | 11.798 | 13.119 | 13.327 | 10.699 | 10.270 | 8.989 | 8.115 | 8.981 | 9.316 | 6.810 | **--** | 1.185 | 1.280 |
| 13. *P. vaccarum* | 8.090 | 6.592 | 12.165 | 12.400 | 10.171 | 10.352 | 10.886 | 9.724 | 9.566 | 9.578 | 11.592 | 12.389 | **1.982** | 0.526 |
| 14. *P. vaccarum** | 7.806 | 6.516 | 12.072 | 11.992 | 9.546 | 9.643 | 10.502 | 9.017 | 8.941 | 9.330 | 11.109 | 12.027 | 3.084 | **0.544** |

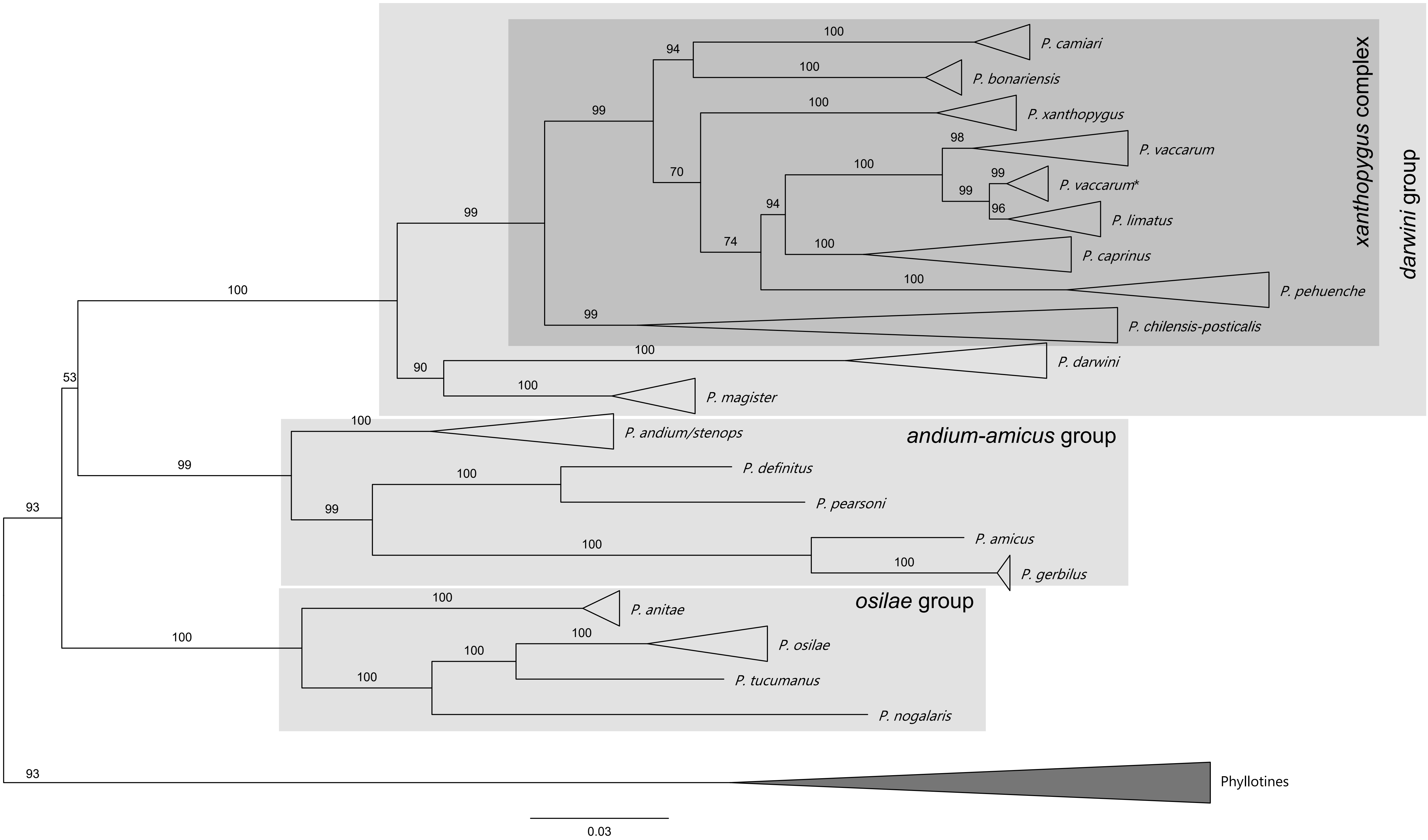


**Supplementary Figure 1.** Phylogenetic relationships of *Phyllotis* species estimated from a ML analysis of 454 cytb gene sequences. ML consensus tree (ln = -7890.962). Supports for internal nodes are shown as ultrafast bootstrap values. Clades at species level are collapsed to simplify visualization of relationships. Specimens in the clade labeled *P. vaccarum** carry cytb haplotypes that is sister to the clase of haplotypes of *P. limatus*, even though whole-genome sequence data confirmed their identity as *P. vaccarum* (Storz et al., 2024).

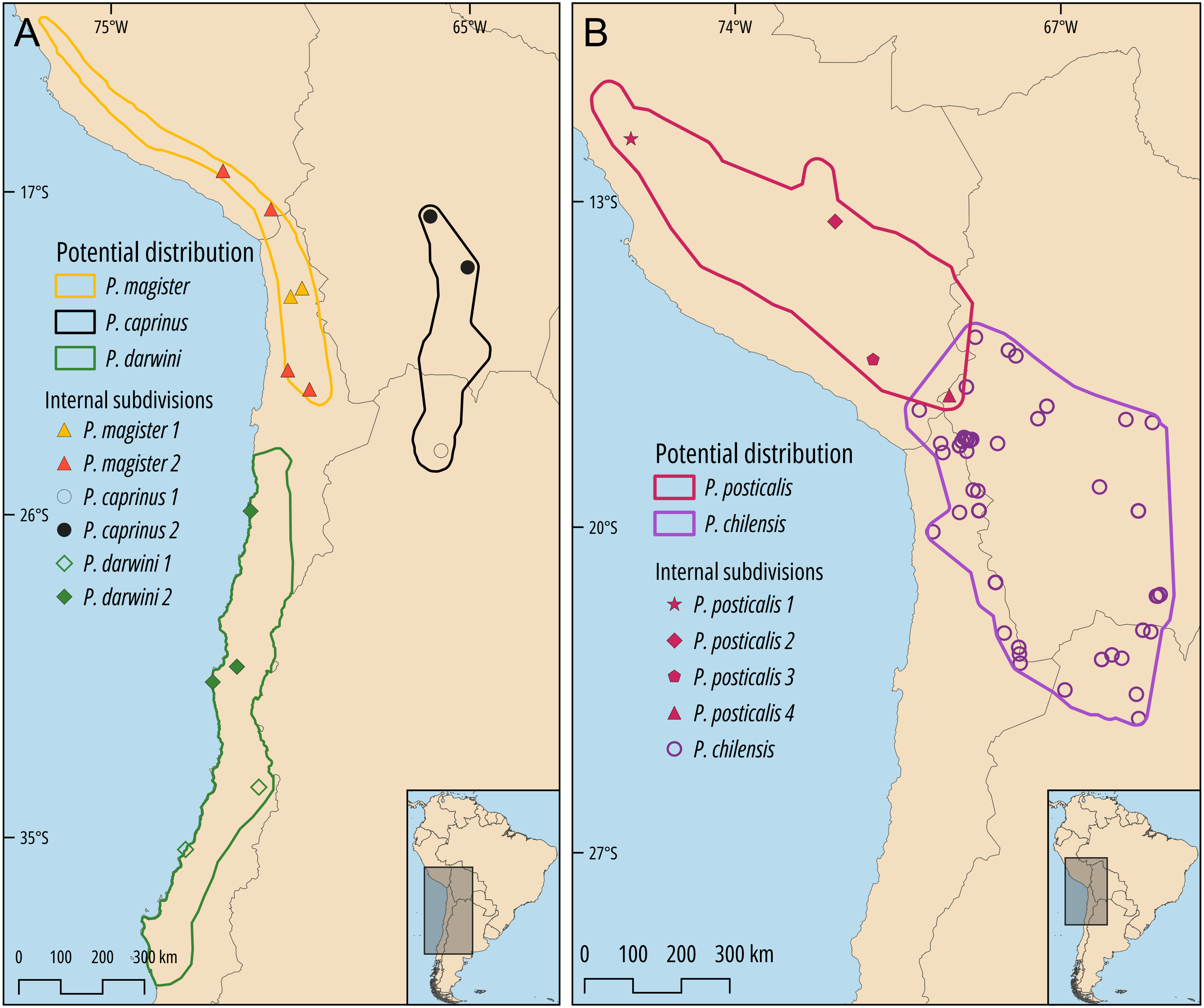


**Supplementary Figure 2.** Geographic distributions of highly divergent, internal subdivisions within several species in the *Phyllotis darwini* group. A) Distribution of representatives of genetically distinct subdivisions within *Phyllotis caprinus*, *P. darwini*, and *P. magister*. B) Distribution of representatives of internal clades of "*P. chilensis-posticalis*". For reasons explained in the text, we regard representatives of the clade with the southernmost distribution as "*P. chilensis*", whereas representatives of the more northern, Peruvian clades are provisionally labelled "*P. posticalis* 1, 2, 3, and 4". Numbers for each clade correspond to their position in the tree shown in Figure 2.
